# Big city, small world: Density, contact rates, and transmission of dengue across Pakistan

**DOI:** 10.1101/018481

**Authors:** M.U.G. Kraemer, T.A. Perkins, D.A.T. Cummings, R. Zakar, S.I. Hay, D.L. Smith, R.C. Reiner

## Abstract

Macroscopic descriptions of populations commonly assume that encounters between individuals are well mixed; i.e., each individual has an equal chance of coming into contact with any other individual. Relaxing this assumption can be challenging though, due to the difficulty of acquiring detailed knowledge about the non-random nature of encounters. Here, we fitted a mathematical model of dengue virus transmission to spatial time series data from Pakistan and compared maximum-likelihood estimates of “mixing parameters” when disaggregating data across an urban-rural gradient. We show that dynamics across this gradient are subject not only to differing transmission intensities but also to differing strengths of nonlinearity due to differences in mixing. We furthermore show that neglecting spatial variation in mixing can lead to substantial underestimates of the level of effort needed to control a pathogen with vaccines or other control efforts. We complement this analysis with relevant contemporary environmental drivers of dengue.

## Introduction

The transmission dynamics of all infectious diseases depend on a few basic but key determinants: the availability of susceptible and infectious hosts, contacts between them, and the potential for transmission upon contact. Susceptibility is shaped primarily by historical patterns of transmission, the natural history of the pathogen, the host’s immune response, and host demography (Grenfell *et al.* 2004). What constitutes an epidemiologically significant contact depends on the pathogen’s mode of transmission (Stoddard *et al.* 2009), and structure in contact patterns can be influenced by transportation networks and the spatial scale of transmission (Brockmann & Helbing 2013), by host heterogeneities such as age (Kilpatrick *et al.* 2006), and dynamically in response to the pathogen’s influence on host behavior (Fenichel *et al.* 2011). Whether transmission actually occurs during contact between susceptible and infectious hosts often depends heavily on environmental conditions (Shaman *et al.* 2010; Gilbert *et al.* 2014; Weiss *et al.* 2014b). Disentangling the relative roles of these factors in driving patterns of disease incidence and prevalence is a difficult but central pursuit in infectious disease epidemiology, and mathematical models that are specific about the biology of how these mechanisms interact represent an indispensible tool in this pursuit (Anderson & May 1991).

The time-series susceptible-infected-recovered (TSIR) model was developed by Finkenstädt & Grenfell (2000) to offer a more accurate and straightforward way to statistically connect mechanistic models of infectious disease transmission with time series data. Among other features, TSIR models readily account for inhomogeneous mixing in a phenomenological way by allowing for nonlinear dependence of rates of contact between susceptible and infectious hosts on their densities. Although this is a simple feature that can be incorporated into any model based on mass-action assumptions – indeed, earlier applications pertained to inhomogeneous mixing in predator-prey systems (Pascual & Levin 1999) – the “mixing parameters” that determine the extent of this nonlinearity have primarily been fit to empirical data in applications of the TSIR model to transmission of measles, cholera, rubella, and dengue (Bjørnstad *et al.* 2002; Koelle & Pascual 2004; Metcalf *et al.* 2011; Reich *et al.* 2013). Applied to discrete-time models such as the TSIR, mixing parameters also have an interpretation as corrections for approximating a truly continuous-time process with a discrete-time model (Liu *et al.* 1987; Glass *et al.* 2003). In no application of the TSIR model to date has the potential for variation in these parameters been assessed, leaving the extent to which inhomogeneity of mixing is different in space and time as an open question in the study of infectious disease dynamics.

There are a number of reasons why mixing might vary in time or space. Seasonal variation in mixing might arise because of travel associated with labor (Bharti *et al.* 2011), religious events (Lessler *et al.* 2014), or vacation (Deville *et al.* 2014). A major driver of flu is associated with the timing of school openings (Salathé *et al.* 2010), social networks (Cauchemez *et al.* 2011), and spatial variation in mixing could arise because of cultural differences and the geographic scales of interest (Brockmann & Helbing 2013; Vazquez-Prokopec *et al.* 2013; Read *et al.* 2014), because of variation in the density and quality of roads (Uchida & Nelson 2008), or because of variation in human densities and myriad factors associated with that (Bjørnstad *et al.* 2002; Simini *et al.* 2012). For vector-borne diseases, variation in mixing is amplified even further by variation in vector densities (Perkins *et al.* 2013), which effectively mediate contact between susceptible and infectious hosts.

Dengue is a mosquito-borne viral disease with a strong potential for spatial variation in mixing (Brady *et al.* 2012; Bhatt *et al.* 2013). The dominant vectors of dengue viruses (*Aedes spp.*) thrive in areas where they are able to associate with humans, as humans provide not only a preferred source of blood but also water containers that the mosquitoes use for egg laying and for larval and pupal development (Morrison *et al.* 2008). Two additional aspects of *Aedes* ecology – limited dispersal distance of the mosquito (Harrington *et al.* 2005) and daytime biting (Akram *et al.* 2009) – imply that human movement should be the primary means by which the viruses spread spatially (Stoddard *et al.* 2009). Indeed, analyses of dengue transmission dynamics at a variety of scales have strongly supported this hypothesis (Allicock *et al.* 2012; Stoddard *et al.* 2013; Bhoomiboonchoo *et al.* 2014; Reiner *et al.* 2014). To the extent that human movement in dense urban environments is more well-mixed than elsewhere, there are likely to be differences in the extent of inhomogeneous mixing in peri-urban and rural areas. This is also presumably the case for directly transmitted pathogens, but with a potentially even stronger discrepancy for dengue due to the urban-rural gradient in mosquito densities.

To assess the potential for spatial variation in the inhomogeneity of mixing as it pertains dengue transmission, we performed an analysis of district-level time series of dengue transmission in the Punjab province of Pakistan using a TSIR model with separate mixing parameters for urban and rural districts. We likewise made estimates of the relationships between density-independent transmission potential and putative drivers thereof, such as temperature, to allow for the relative roles of extrinsic and intrinsic factors to be teased apart. Finally, we performed mathematical analyses of the fitted model to assess the significance of spatial variation in mixing inhomogeneity for how time series data are interpreted and used to guide control efforts.

## Material and Methods

We obtained daily dengue case data aggregated at hospital level from Punjab province provided by the Health Department Punjab, Pakistan between 2011 and 2014. In total, 41,300 suspected and confirmed dengue cases were reported in 109 hospitals. All hospitals were subsequently geo-located using ‘Google maps’ (http://www.maps.google.com). Hospitals that could not be identified were removed from the database. 21,182 cases alone were reported from the year 2011, which affected the almost the entire province. Many more cases occurred in Lahore (35,348) compared to all other districts (5,952) (Table 1). A breakdown per year and each province is provided in Table S1 and additional information about collection can be found in the supplementary appendix.

**Table 1:** Reported cases by year in Lahore and all other districts.

|  | 2011 | 2012 | 2013 | 2014 | Total |
| --- | --- | --- | --- | --- | --- |
| Lahore | 18,020 | 4,013 | 11,516 | 1,799 | 35,348 |
| Other | 3,162 | 500 | 2,356 | 2,790 | 5,952 |

### Covariate selection and processing

Environmental conditions are instrumental in defining the risk of transmission of dengue (Bhatt *et al.* 2013). Transmission is limited by the availability of a competent disease vector. Due to a lack of resources and political instability no comprehensive nation wide entomological surveys have been performed in Pakistan. Therefore we use a probabilistic model to infer the probability of occurrence of *Ae. aegypti* and *Ae. albopictus* in Pakistan derived from a globally comprehensive dataset containing more than 20,000 records for each species (Figure 1a and b). Such model outputs have proven useful in identifying areas of risk of transmission of dengue as well as malaria (Gething *et al.* 2011; Sinka *et al.* 2012; Bhatt *et al.* 2013). Other important environmental conditions defining the risk of transmission of dengue are temperature, water availability, and vegetation cover (Messina *et al.* 2015). To account for such variation, raster layers of daytime land surface temperature (LST) were processed from the MOD11A2 satellite, gap-filled to remove missing values, and then averaged to a monthly temporal resolution for all four years (Weiss *et al.* 2015). The density of vegetation coverage has been shown to be associated with vector abundance (Eisen & Lozano-Fuentes 2009). Moreover, vegetation indices are useful proxies for precipitation and may be used to derive the presence of standing water buckets that are habitats for the *Aedes* mosquitos (Maciel-de-Freitas *et al.* 2007). The same method was again applied to derive the Enhanced Vegetation Index (EVI) from the MOD11A2 satellite to produce 16 day and monthly pixel based estimates for 2011-2014 (Figure 1g) (Weiss *et al.* 2014a). Due to the inherent delay between rainfall and daily temperature influencing mosquito population dynamics and those mosquitoes contributing to an increase in DENV transmission, we consider both, the influence of the current temperature, vegetation indices and precipitation, data on current transmission as well as the values of those covariates the time step before (Figure 1f).

**Figure 1:**
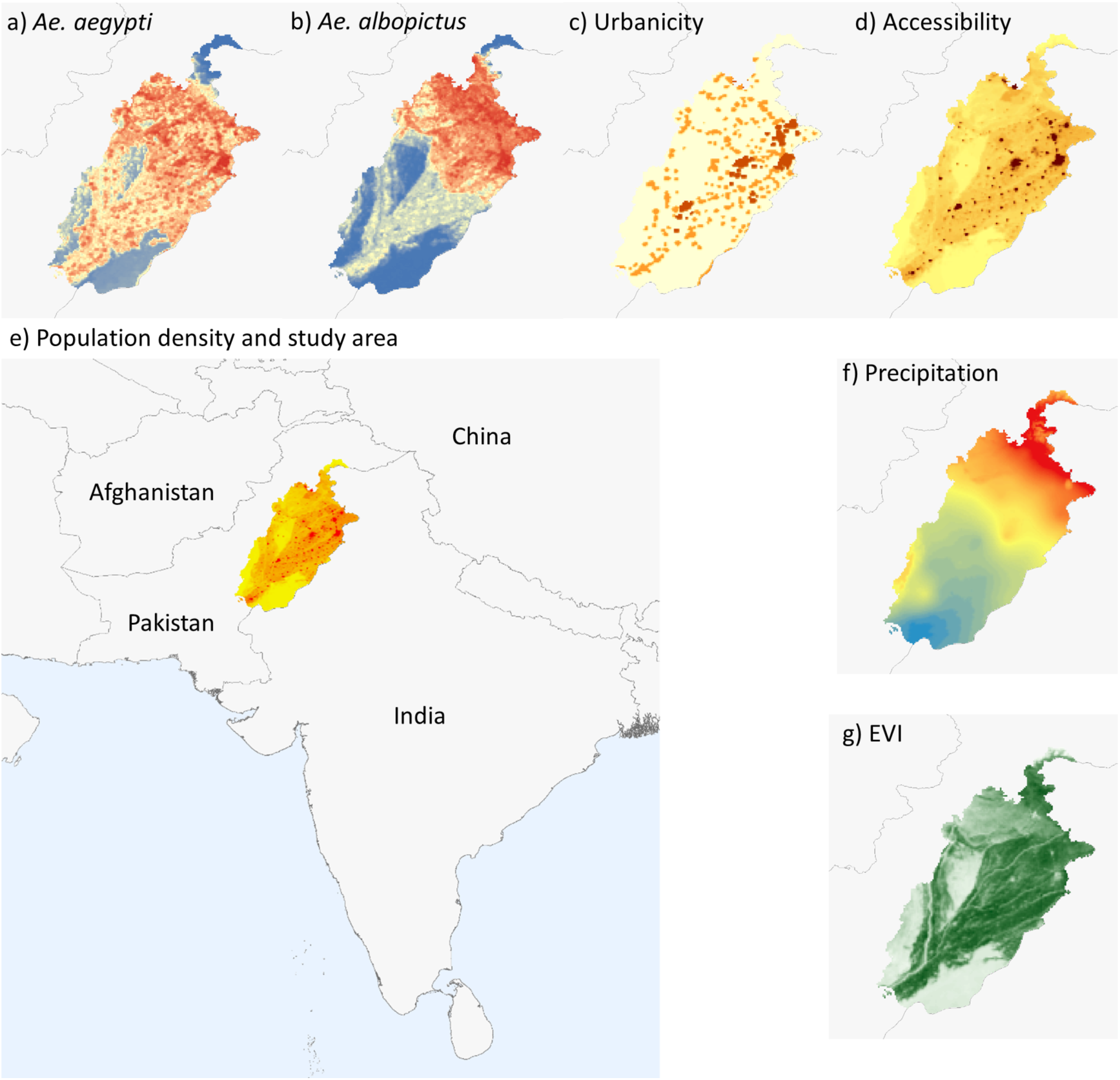
Covariates used in this study to derive environmental drivers of transmission. *Aedes aegypti* probability of occurrence (a); *Ae. albopictus* probability of occurrence (b); Urbanicity (c); Weighted urban accessibility (d); Population density and study area (e); Precipitation (f); Enhanced Vegetation Index (EVI) mean (g).

We used population count estimates on a 100m resolution that were subsequently aggregated up to match all other raster layers to a 5 km × 5 km resolution for the year 2015 (<u>http://www.worldpop.org</u>) (Figure 1e). In a follow-up analysis to our primary investigation into the climatological drivers of dengue transmission we included several density-based covariates. We derived a weighted accessibility metric that includes both, population density and urban accessibility, a metric commonly used to derive relative movement patterns (Uchida & Nelson 2008; Tatem *et al.* 2012). This map shows a friction surface, i.e. the time needed to travel through a specific pixel (Figure 1d). We also used an urban, peri-urban and rural classification scheme to quantify patterns of urbanicity based on a globally available grid (Center for International Earth Science Information Network (CIESIN) 2010) (Figure 1c). All covariates and case data were aggregated and averaged (where appropriate) to a district level.

### Model

Following Finkenstädt & Grenfell (2000), we assume a general transmission model of

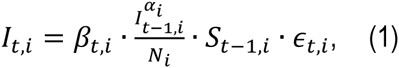

where *I*_*t,i*_ is the number of infected and infectious individuals and *S*_*t,i*_ the number of susceptible individuals, at time *t* in district *i, N*_*i*_ is the population of district *i*, and *β*_*t,i*_ is the covariate driven contact rate. The mixing parameter for the *i*th district is given by *α*_*i*_; when *α*_*i*_ is equal to 1, the population has homonegeous mixing where values less than one can either indicate inhomogeneous mixing or a need to correct for the discretization of the continuous-time process. Finally, the error terms *∈_t,i_* are assumed to be independent and identically log-normally distributed random variables. For more details please see SI.

### Model selection

The term *β*_*t,i*_ is fit using generalized additive model regression (GAM) (Hastie & Tibshirani 1990; Dominici *et al.* 2002; Wood 2011). The time-varying climatological covariates are all fit as a smooth spline, while all other covariates enter *β*_*t,i*_ linearly. Additionally, unexplained seasonal variation is accounted for using a 12-month periodic smooth spline.

Model selection was performed using backwards selection. Two base models were investigated. First, a climate only model was created using only the climatological and climate suitability covariates. Second, a “full” model using the density-dependent covariates as well as the climatological covariates were combined into a single model which was then subjected to backwards selection. For both models the mixing coefficient was initially set equal for each district and once a final model was arrived upon, the mixing coefficient for Lahore was allowed to vary separately from the other coefficients. All model fitting was conducted using R (R Core Team 2014) and the “mgcv” package (Wood 2011). Models are fit by maximizing the restricted maximum likelihood (Patterson & Thompson 1971) to reduce bias and over-fitting of the smooth splines. The model source code and processing of covariates will be made available in line with previous projects (Pigott & Kraemer 2014).

### Model analysis

To explore the potential significance of spatial variation in mixing parameters, we conducted an analysis to probe the inherent mathematical tradeoff between the mixing parameter *α* and the density-independent transmission coefficient *β*. Specifically, to answer the question, what difference in local transmission would be necessary to account for a difference in mixing while achieving identical transmission dynamics. To explore this, we used eqn. (1) to establish:

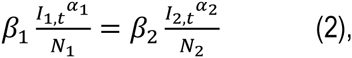

from which we obtained

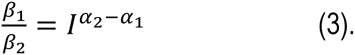

We then examined how variation in *l* and *α*_2_ − *α*_1_ affected the left hand side of eqn. (3) and likewise the critical proportion of the population to control in order to effect pathogen elimination, which, under our model, is 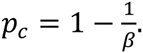.

## Results

### Description of case distribution

The majority of cases are clustered in Lahore, the capital of Punjab province. Ongoing transmission seems to be focal in three (Vehari, Rawlpindi and Lahore) districts and spread through infective sparks to smaller more rural provinces. In early investigations, all hospitals that reported dengue cases are located in areas with high EVI values indicating a clear environmental signal (Figure 1b).

### Model selection

To disentangle the different aspects of dengue dynamics and their drivers we used a model containing only the climatological covariates and performing backwards model selection until each covariate in the model was significant at a 5% level resulted in a model that explained 76.9% of the deviance and whose adjusted R-squared was 0.746. Amongst the yearly-averaged covariates, EVI and precipitation remained in the model as well as the derived *Ae. albopictus* range map (p-values of 8.21 × 10^-4^, 0.01, and 3.9 × 10^-5^ respectively). Interestingly, if the derived *Ae. aegypti* map is substituted for the *Ae. albopictus* map, the deviance explained increases slightly to 76.8%. For climatological covariates that were fit as smooth splines, temperature, lagged temperature and EVI remained in the model (Figure 2, p-values of 0.010, 0.030 and 0.030 with effective degrees of freedom 7.55, 5.47, and 1.83, respectively). There was a significant amount of periodic variation unexplained by the climatological covariates alone and the ‘seasonality’ covariate remained (Figure 2, p-value = 0.0034). The estimated median values for R_0_ per district are clustered around two (mean = 2.1), their geographical distribution indicates a clear trend towards districts with higher population (Figure 3). Finally, the mixing coefficient was significantly lower than 1 (*α* = 0.69, 95% CI = (0.614, 0.771), p-value = 1.6 × 10^−14^).

**Figure 2:**
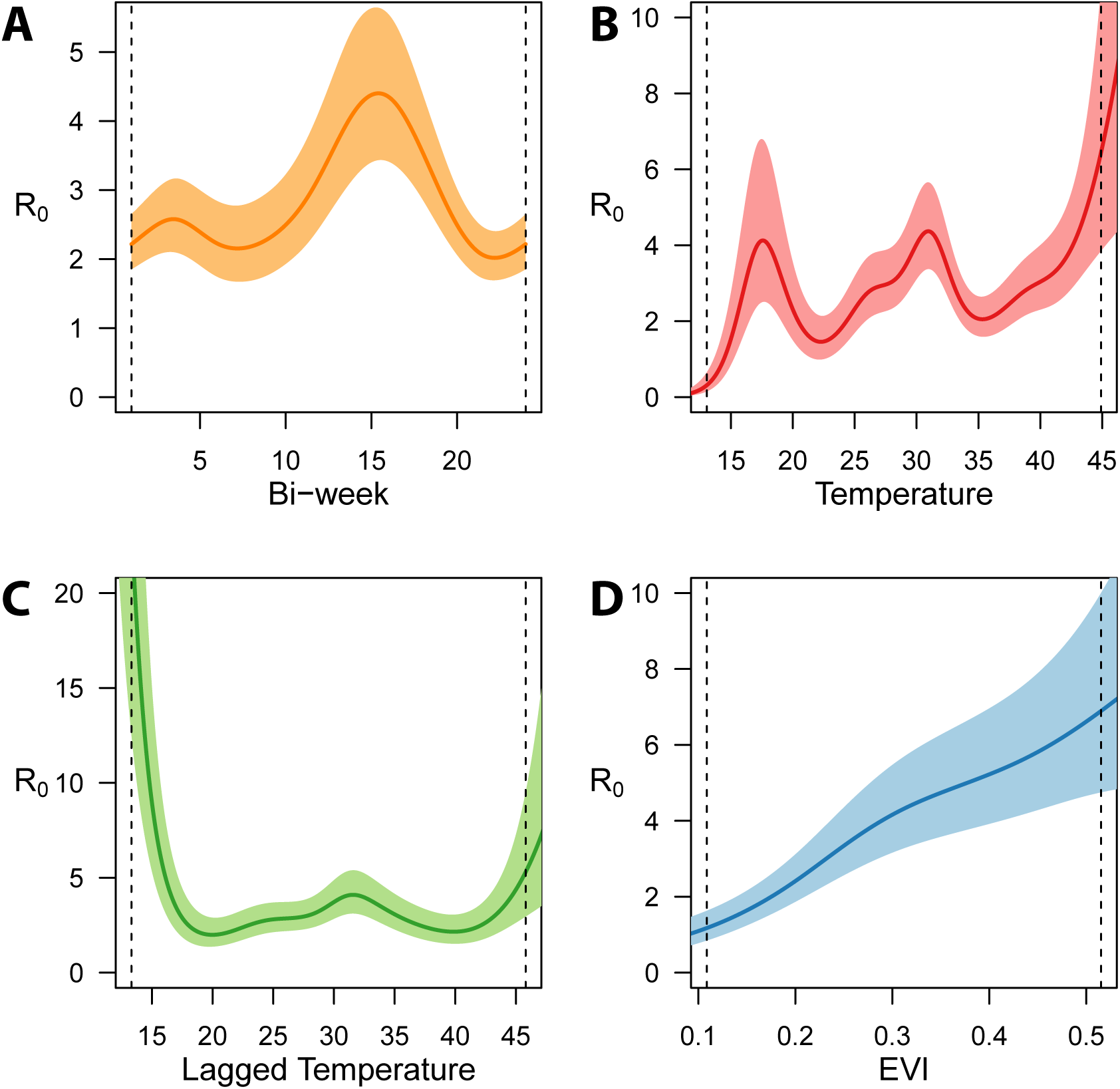
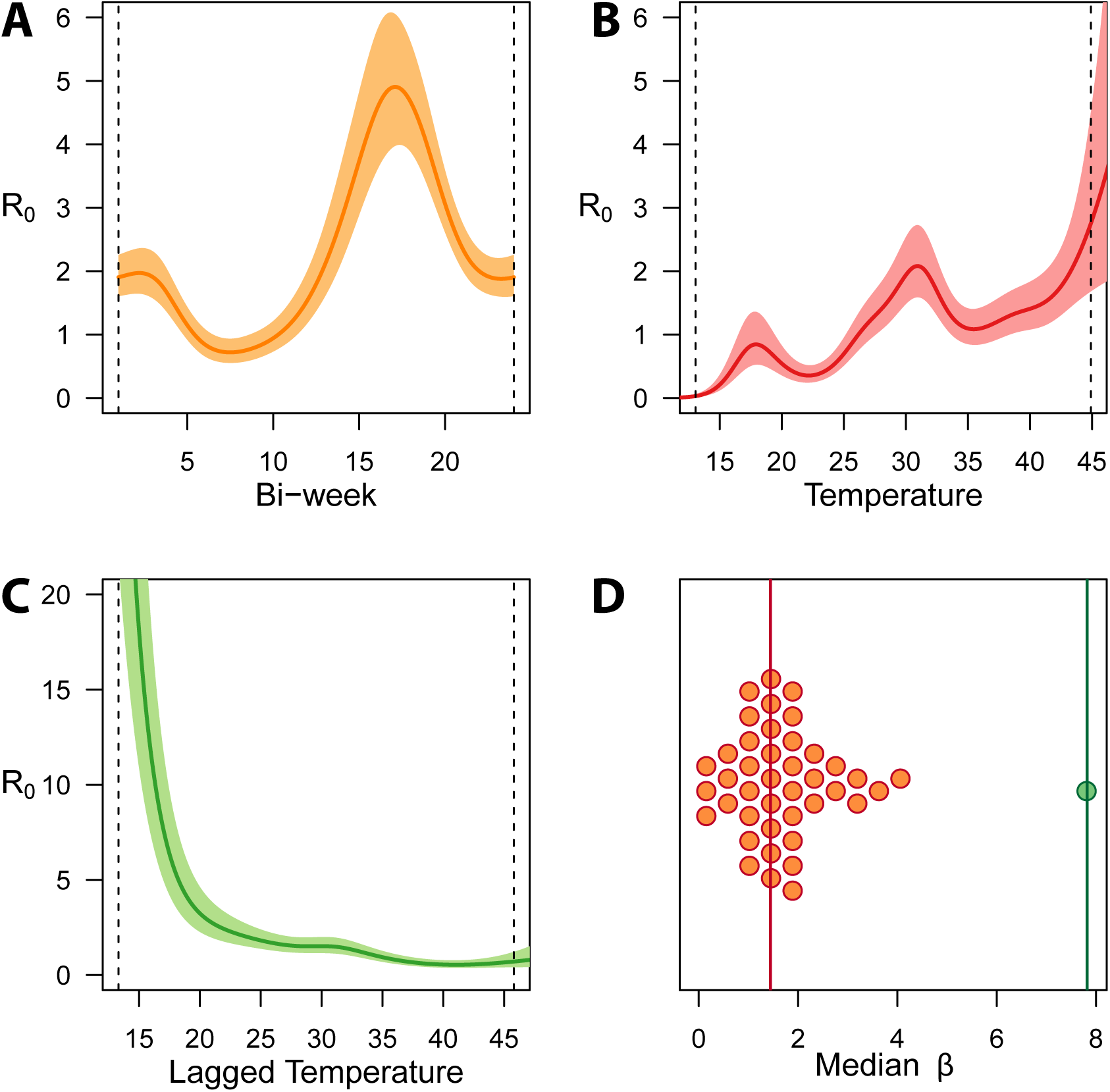
Model outputs using a backwards model selection procedure in the model using climatological variables (a, A-D), and including the density dependent variables (b, A-D).

**Figure 3:**
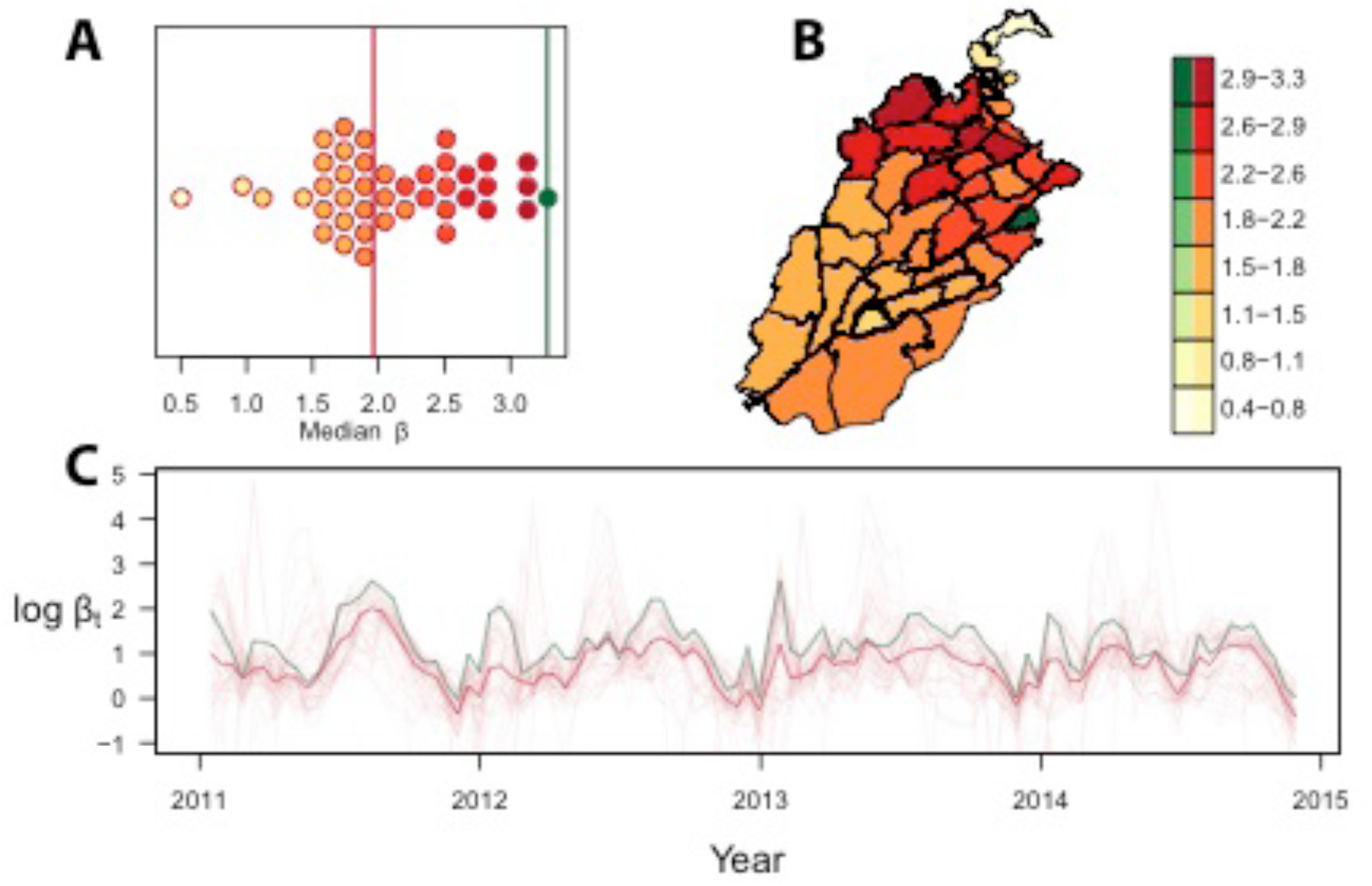
Average distribution of R_0_ with green representing Lahore versus red all other districts (A), their geographical distribution (B), and over time in which the green line again is representing Lahore versus the red line representing all other districts (C).

To understand these differences the final model was then compared to a nested model where the coefficient for Lahore was allowed to vary independently of all other districts. Deviance explained increased to 77% and adjusted R-squared increased to 0.753. Further, the mixing coefficient for Lahore (*α* = 0.74) was significantly larger than the mixing coefficient for the other districts (*α* = 0.59, p-value=0.0068) (Figure S1). The median R_0_ for Lahore was estimated at 3.28, the highest for all districts.

In assess how far the variation of mixing coefficients can be explained by other covariates we consider the possibility that movement accounts for the differences in the mixing coefficient between Lahore and the other districts. The density-dependent covariates (described earlier) were then added to the full model and backwards selection was repeated. The resulting model explained 78.6% of the variance, had an adjusted R-squared of 0.763 and is superior to the final climatological model based on AIC (699.23 versus 714.83). Yearly averaged EVI, NDVI and precipitation were all significant (p-values of 8.7 × 10^-5^, 0.00024, and 0.00028, respectively). Again, the derived *Ae. albopictus* map was significant (p-value = 0.008163). For climatological covariates fit as smooth splines, only temperature and lagged temperature were found significant (Figure 2b, p-value of 4.0 × 10^-5^ and 0.0013, and effective degrees of freedom 7.61 and 4.81, respectively), and there was still a significant ‘seasonality’ (Figure 2b, p-value of 4.0 × 10^-7^, effective degrees of freedom 4.48). The mixing coefficient was fit at *α* = 0.58, barely lower than the mixing coefficient for non-Lahore districts in the climatological model. The estimated median R_0_ again clustered around 2 (mean = 1.8), and again the R_0_ for Lahore was largest, but in this model it was considerably larger than in the climatological model (Lahore R_0_= 7.82, Figure 2b). Full details of the model parameters are shown in Table S2-S5.

Two of the density-dependent covariates remained in the model: the urban map (p-value = 0.01) and the weighted access map (p-value= 3 × 10^-5^). When the nested model that allows Lahore’s mixing coefficient to vary was fit, there was no significant difference in the two mixing coefficients (p-value≈1).

### Model analysis

Given a difference in estimates of the mixing parameters between Lahore and elsewhere of 0.15, we analyzed eqn. (3) to assess the bias in estimates of the transmission coefficient that would result from ignoring this extent of variation in the inhomogeneity of mixing displayed between two areas. For the purpose of *ceteris paribus* comparisons, we assumed equal force of infection but varied it across several orders of magnitude. Depending on the order of magnitude, estimates of transmission coefficients made if overestimating the mixing parameter by 0.15 could easily result in a two- to three-fold underestimate in the transmission coefficient (Figure 4). For realistic ranges of the transmission coefficient for dengue, and more generally the basic reproductive number R_0,_ this extent of underestimation of R_0_ could lead to underestimating the critical proportion of the population to which vaccines or other control measures must be applied by 20-30% (Figure 5).

**Figure 4:**
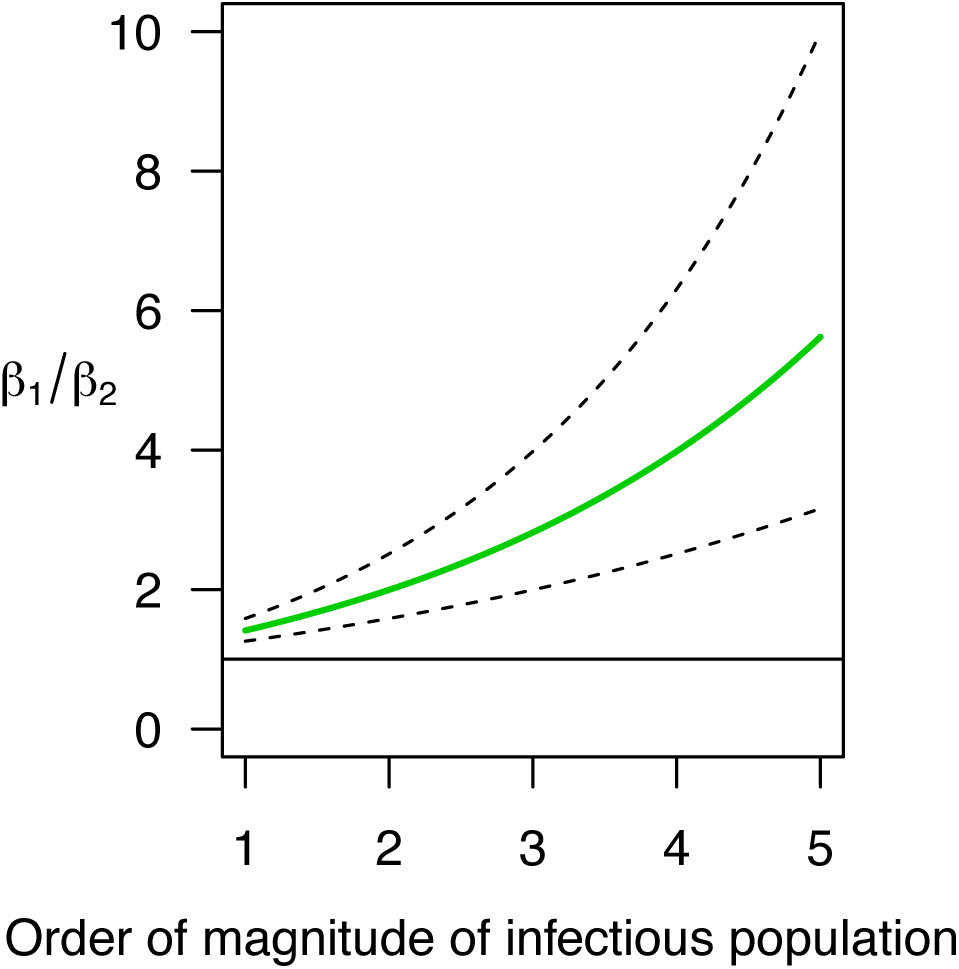
Ratio of betas (R_0_) assuming equal force of infection and a difference in *α*_2_ – *α*_1_ of 0.2, 0.15 (green), and 0.1, form top to bottom. The straight line indicates a ratio of 1.

**Figure 5:**
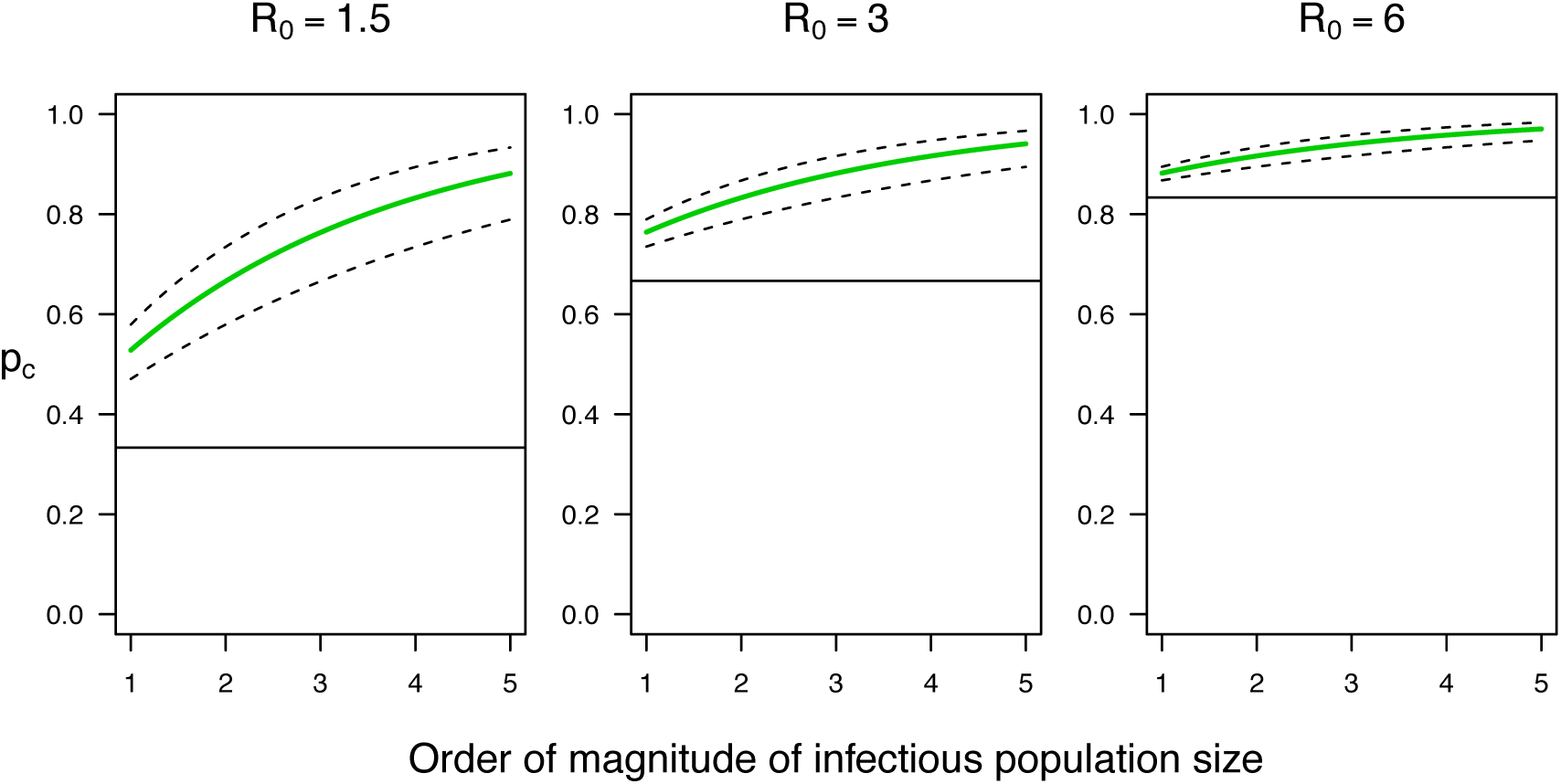
Critical proportion of the population to control in population 2 as a function of R_0_ in population 1, the order of magnitude of the infectious numbers in each population, and a difference in *α*_2_ – *α*_1_ of 0.1, 0.15 (green), and 0.2. The straight line indicates the critical proportion assuming the *α* in each population are equal.

## Discussion

Our results point to considerable spatial heterogeneity in the extent of mixing and strength of an associated nonlinearity in transmission along an urban-rural gradient. This regional variability in mixing has direct implications for estimates of the basic reproductive number of dengue in our study region and elsewhere. Although the potential for such bias in estimates of the basic reproductive number has been shown in a theoretical context (Hu *et al.* 2013; Perkins *et al.* 2013), we provide quantitative estimates of the extent of this problem in interfacing models with a rich spatio-temporal data set. Such analyses have implications for estimates of population-level parameters not only for dengue but also for other infectious diseases (Bartlett 1960; Keeling & Grenfell 1997; Bjørnstad *et al.* 2002; Smith *et al.* 2002; Keeling & Eames 2005; Kiss *et al.* 2008) and possibly even more broadly in ecology.

We revealed significant differences in mixing components between urban and rural settings and found that a population-weighted urban accessibility metric was able to absorb differences in mixing between these settings indicating that this specific covariate accounts for aspects influencing mixing. Mixing is presumably influenced directly by human behavior and has been shown to be highly unpredictable, largely dependent on the local context and the spatial and temporal scale (Yang *et al.* 2014). In this study however we could show that the density-dependent covariate selected was able to capture the influence of these key encounters on a district level. Once differences in mixing were accounted for, estimated R_0_ values indicated considerably larger differences between transmission potential in Lahore versus all other districts. Synchronizing more accurate geo-referenced data would allow to assess the importance of spatial scale on the relationship between “mixing parameters” and urban accessibility (Perkins *et al.* 2013; Mills & Riley 2014). In the case of dengue this again has been limited by the availability of high resolution data (Ruberto *et al.* 2015). Complementing this analysis with measurement of direct social contact patterns could be important to explore this relationship in even more detail (Bauch & Galvani 2013; Vazquez-Prokopec *et al.* 2013; Heesterbeek *et al.* 2015) and could be informed by mathematical models that explored this relationship previously for other diseases (Bjørnstad *et al.* 2002; Reiner *et al.* 2012). Another encouraging finding is that large-scale mosquito suitability surfaces help capture the environmental determinants of dengue transmission (Bhatt *et al.* 2013).

Intervention strategies are contingent on both understanding key environmental drivers of transmission and the dynamics of ongoing human-to-human transmission, particularly in outbreaks situations (Perkins *et al.* 2015). Environmental drivers such as seasonal fluctuations in rainfall, temperature, vegetation coverage or mosquito abundance will help guide surveillance and control efforts targeted mostly towards the ecological aspects of mosquito dispersal (Johansson 2015). Once infection occurs much debate has been focused around optimizing intervention strategies to reduce disease incidence, which is largely determined by R_0._ The presented framework shows that the interaction between mixing parameters and force of infection has potentially large implications for optimizing targeted intervention, particularly in countries where resources are scarce (Cesare *et al.* 2015). This in fact is even more important in areas of low infection where transmission seems to be more focal (Salje *et al.* 2012). Again however importance needs to be focused on the spatial and temporal resolution of appropriate intervention strategies and the respective effect of the selected covariates and model parameters (Mills & Riley 2014). Empirical understanding, however, on which spatial resolution is most appropriate to carry out large-scale vector-borne disease interventions remains unknown.

Once transmission has occurred in one place, understanding not only spatial heterogeneities of transmission dynamics but their subsequent spread in mechanistic stochastic models as shown for measles would help to empirically determine the propagation of the disease (Grenfell *et al.* 2001). The spatial spread dynamics have been given considerable interest globally as the risk of importation of dengue into yet endemic areas continentally and internationally is increasing with travel and trade (Schaffner & Mathis 2014). Exploration of the case data in Pakistan suggests that spread happens along major transport routes from Lahore to Karachi and north to Rawalpindi. Using results presented here on mixing components and environmental drivers will help pinpoint areas of major risk of importation more accurately especially in the case of recurring epidemics. Using the fitted relationships of the environmental drivers of transmission and R_0_ will enable future analyses and comparisons between diseases and geographic regions. In this context it will be instrumental to integrate a variety of movement and social network models with the evidence presented here to infer more accurately how the geographical spread of dengue is determined.

To allow for comparison of these results in a broader context and across diseases, possibly even in outbreak situations and in real time, it is essential to make data widely available by open access (Heesterbeek *et al.* 2015). Moreover, the complexity of infectious disease dynamics is not fully understood and we are limited by computational capacities to fully account for stochasticity and nonlinearity.

## Author contributions

Conceived and designed the experiment: MUGK, RCR, TAP, DLS. Performed the experiments: RCR, TAP, MUGK. Analysed the data: RCR, MUGK. Edited the manuscript: DLS, SIH, DATC, ZR. Wrote the paper: MUGK, RCR, TAP.

## Acknowledgements and funding

The authors would like to thank the WHO Punjab office, Health Department Punjab, and PID for providing the epidemiological data, and all participants that helped collect the data. MUGK acknowledges funding from the German Academic Exchange Service (DAAD). TAP, SIH, RCR acknowledge the support from the RAPIDD program of the Science & Technology Directorate, Department of Homeland Security, and the Fogarty International Center, National Institutes of Health. SIH is funded by a Senior Research Fellowship from the Wellcome Trust (#095066) and a grant from the Bill & Melinda Gates Foundation (#OPP1093011). Funders had no role in study design, data collection and analysis, decision to publish, or preparation of the manuscript.

---

**Table S1:**
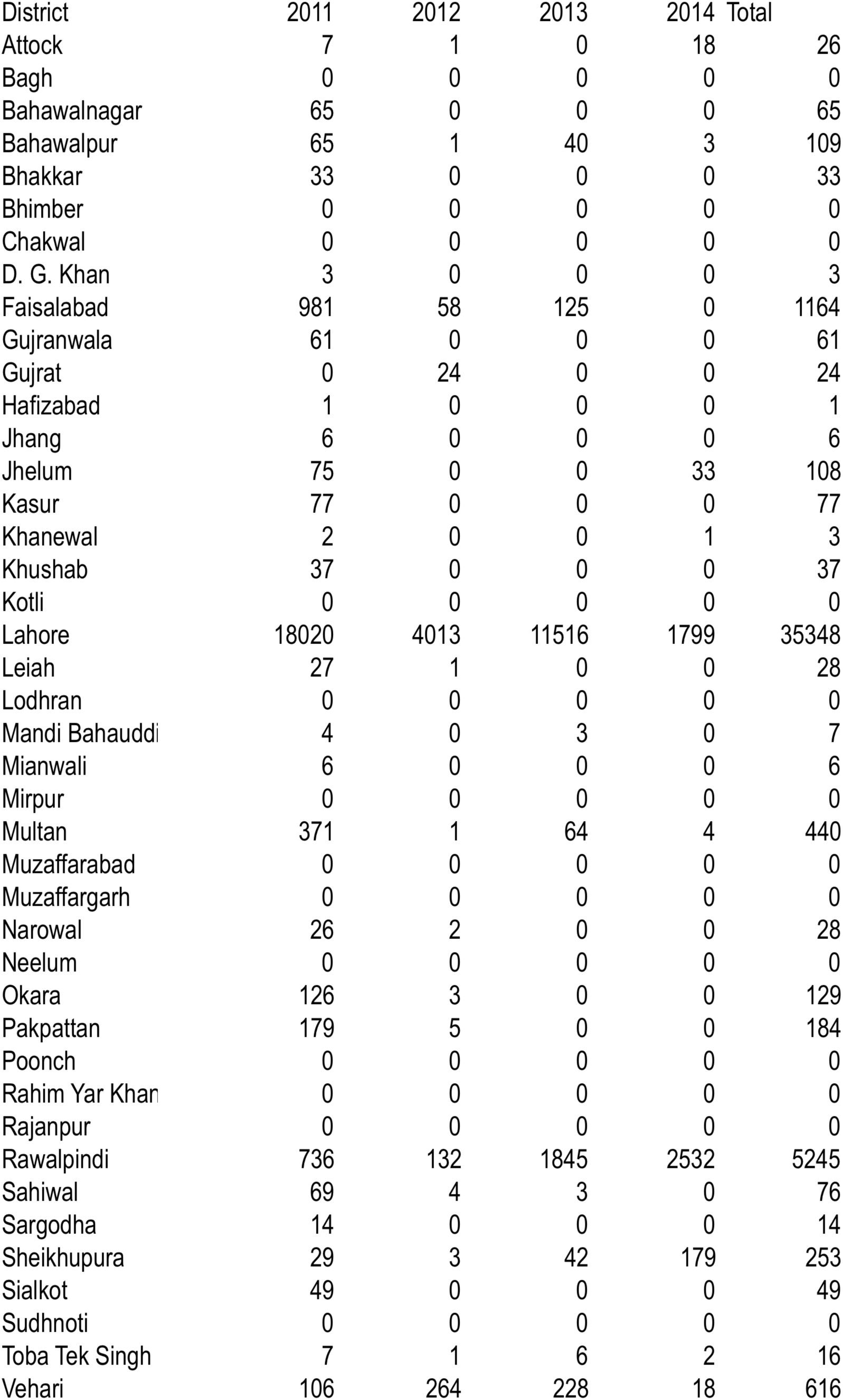
Reported case numbers per year for each district of the study region, Punjab Province, Pakistan.

**Table S2:**
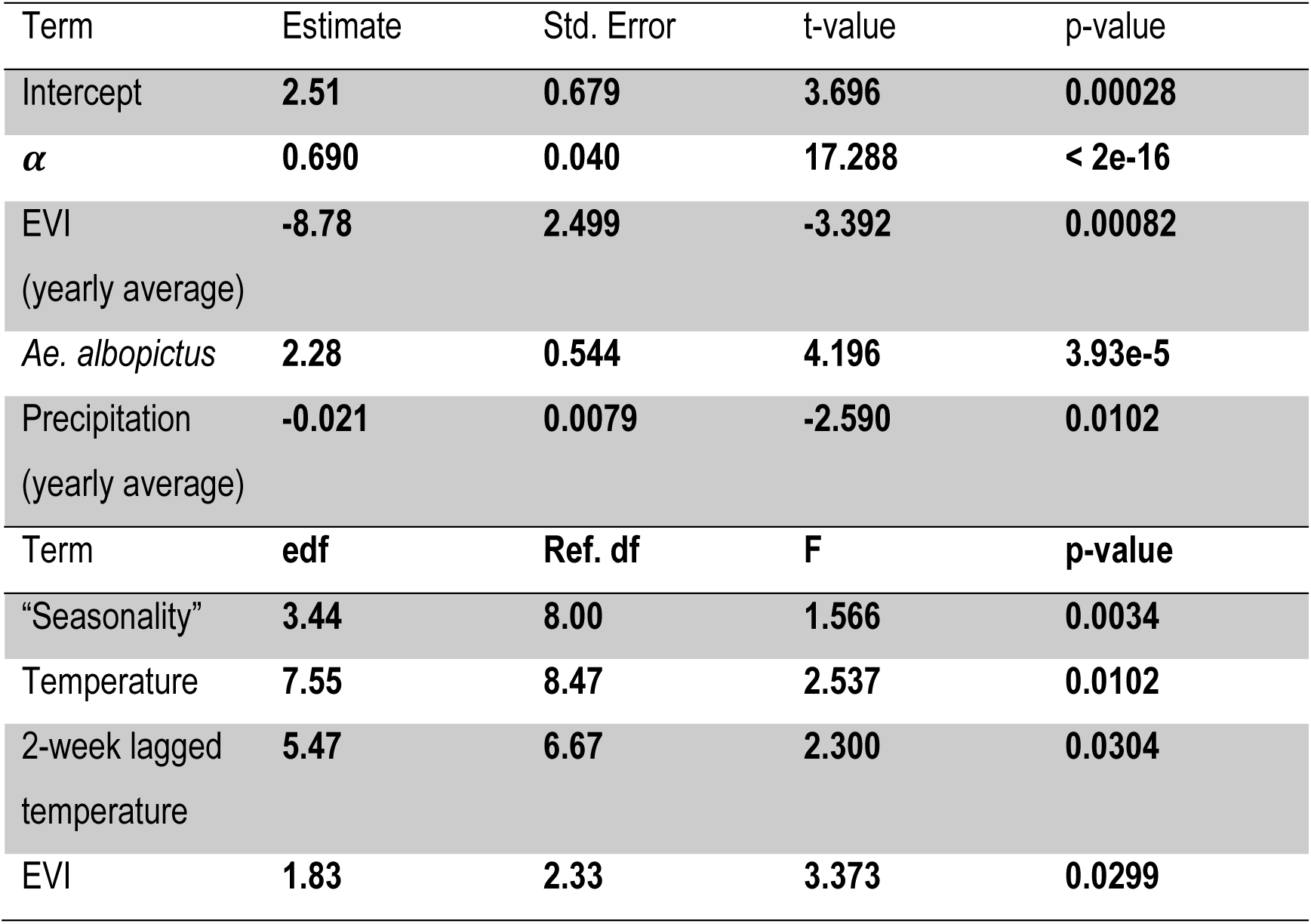
Environmental model: no variation in *α*.

**Table S3:**
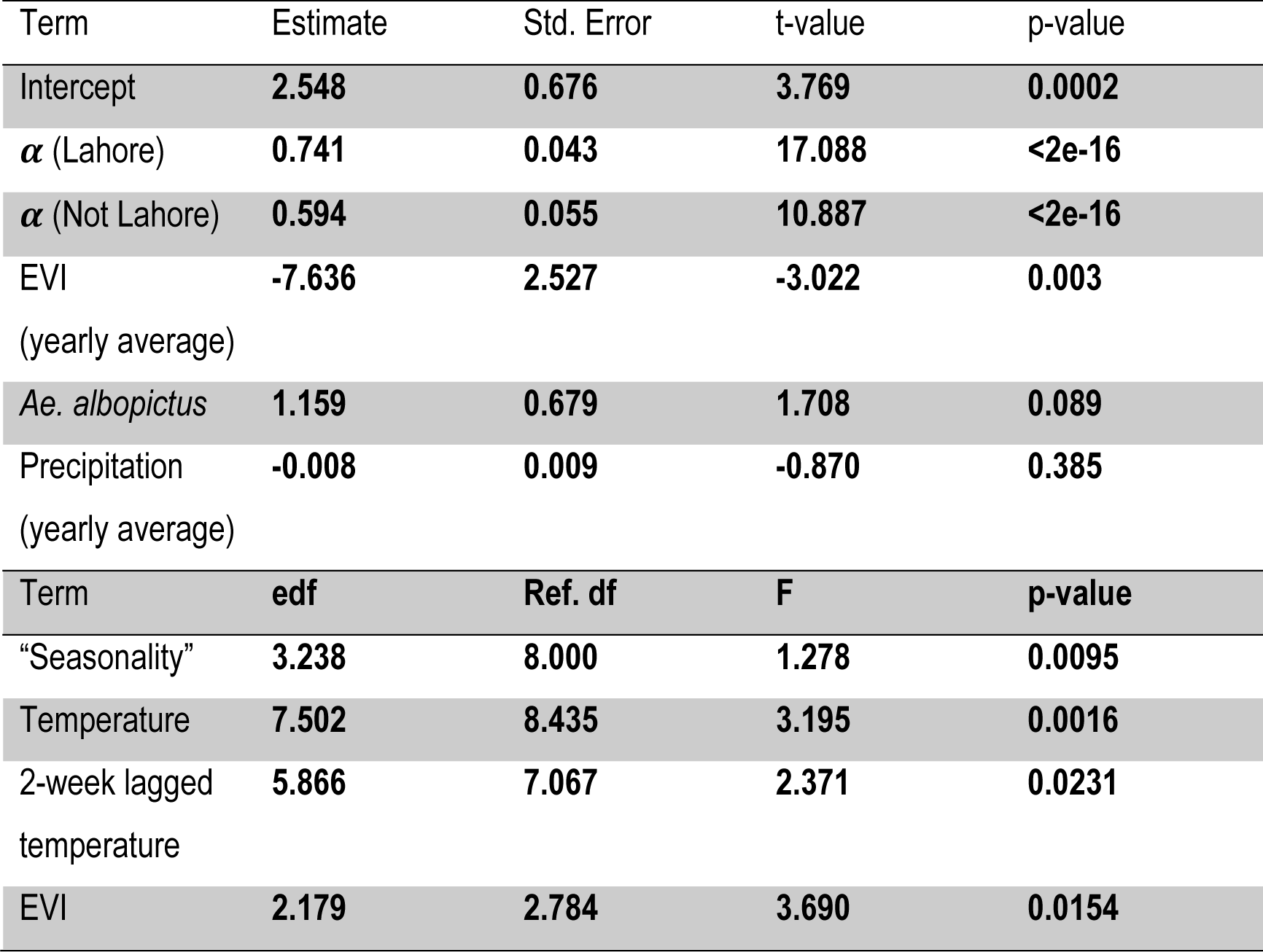
Environmental model: *α* in Lahore differs from other districts.

**Table S4:** Full model: no variation in *α*.

| Term | Estimate | Std. Error | t-value | p-value |
| --- | --- | --- | --- | --- |
| Intercept | -34.83 | 9.803 | -3.553 | 0.00047 |
| $\alpha$ | 0.578 | 0.0451 | 12.824 | <2e-16 |
| EVI<br>(yearly average) | -60.12 | 15.04 | -3.998 | 8.71e-5 |
| NDVI<br>(yearly average) | 0.0383 | 0.0103 | 3.737 | 0.00024 |
| *Ae. albopictus* | 2.745 | 1.028 | 2.669 | 0.00826 |
| Weighted<br>Access | 2.742e-5 | 6.464e-6 | 4.242 | 3.26e-5 |
| Urbanicity | -2.468 | 0.9494 | -2.600 | 0.00776 |
| Precipitation<br>(yearly average) | -0.0684 | 0.0185 | -3.697 | 0.00028 |
| Term | edf | Ref. df | F | p-value |
| "Seasonality" | 4.480 | 8.000 | 4.173 | 3.97e-7 |
| Temperature | 7.607 | 8.504 | 4.396 | 4.14e-5 |
| 2-week lagged<br>temperature | 4.823 | 5.981 | 3.793 | 0.0013 |

**Table S5:** Full model: *α* in Lahore differs from other districts.

| Term | Estimate | Std. Error | t-value | p-value |
| --- | --- | --- | --- | --- |
| Intercept | -35.92 | 9.93 | -3.618 | 0.00037 |
| $\alpha$ (Lahore) | 0.612 | 0.0643 | 8.510 | <2e-16 |
| $\alpha$ (Not Lahore) | 0.555 | 0.0566 | 9.802 | <2e-16 |
| EVI<br>(yearly average) | -61.59 | 15.19 | -4.055 | 6.95e-5 |
| NDVI<br>(yearly average) | 0.0395 | 0.010 | 3.8 | 0.00019 |
| *Ae. albopictus* | 2.81 | 1.033 | 2.721 | 0.0070 |
| Weighted<br>Access | 2.42e-5 | 7.86e-6 | 3.072 | 0.0024 |
| Urbanicity | -2.33 | 0.9697 | -2.403 | 0.0171 |
| Precipitation<br>(yearly average) | -0.069 | 0.0186 | -3.733 | 0.00024 |
| Term | edf | Ref. df | F | p-value |
| "Seasonality" | 4.350 | 8.000 | 3.817 | 1.31e-6 |
| Temperature | 7.620 | 8.511 | 4.335 | 4.97e-5 |
| 2-week lagged<br>temperature | 4.764 | 5.916 | 3.680 | 0.00178 |

**Figure S1:**
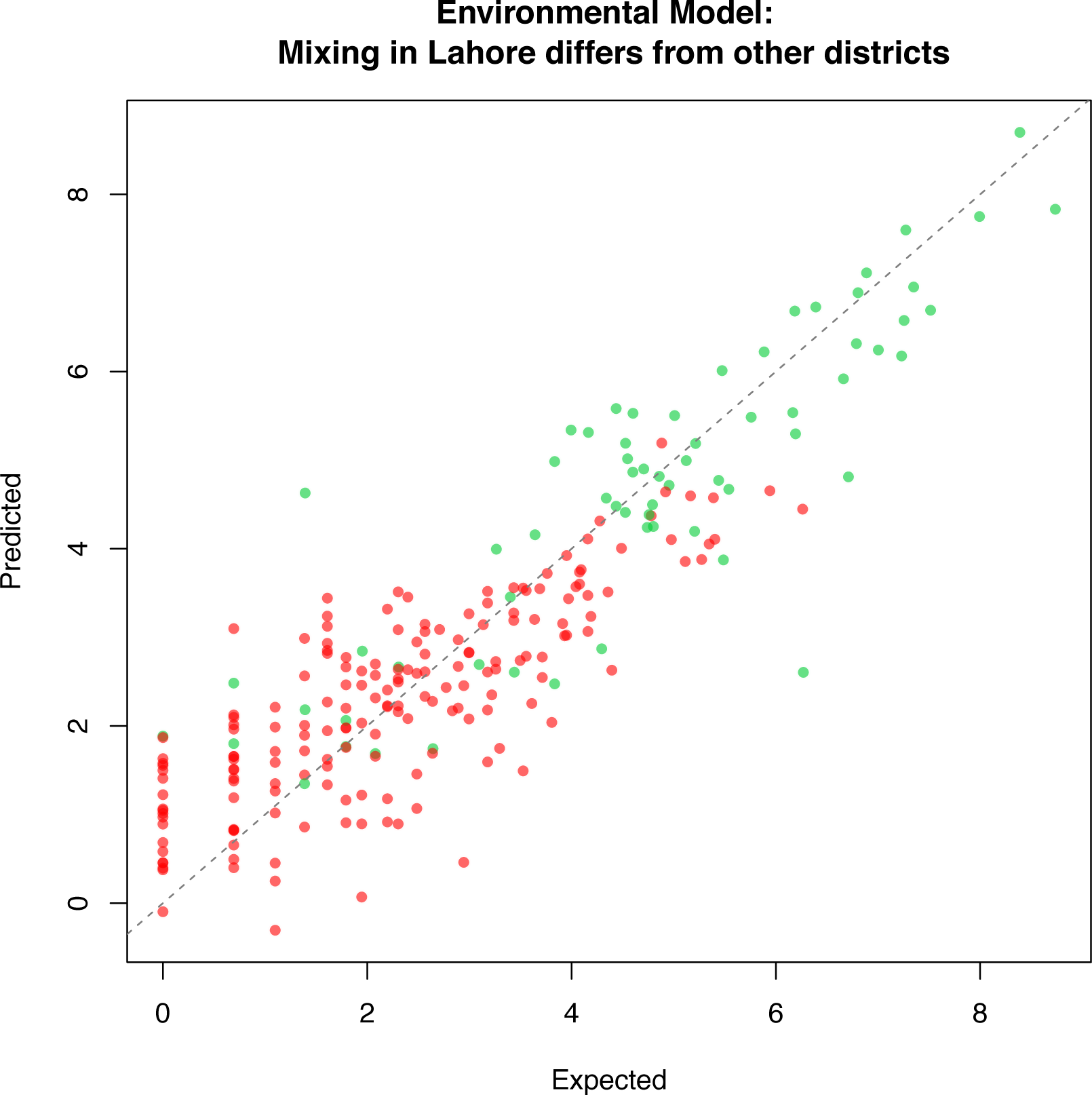
Predicted versus expected values for *α* in Lahore (green) and all other districts (red) for the environmental model.

## Additional information about collection of epidemiological data

Secondary data from hospital records were used from Punjab province in Pakistan. The data was initially collected by Punjab Health Department as part of the dengue prevention and eradication program. For ensuring accurate reporting from the health facilities, Punjab Health Department used the following three procedures: (i) Clinical case reporting, (ii) Lab case reporting, and (iii) Case management. All health facilities were liable to record the data and share them with Punjab Information Technology Board (PITB) within 24 hours. For clinical case reporting, as per Dengue Expert Advisory Group (DEAG), guidelines, dengue suspects, probable, and confirmed cases needed to be correctly entered on the PITB dashboard within 24 hours of admission. For lab case reporting, all private sector labs must send reports of positive dengue cases in the line list format to the respective Executive District Officer Health (EDOH) for online entry on the dashboard, again within 24 hours. For case management, the healthcare facilities are liable to mange dengue cases regularly in specified Dengue care units, OPDs, emergency units, or wards.

